# K-mer analysis of long-read alignment pileups for structural variant genotyping

**DOI:** 10.1101/2024.10.22.619642

**Authors:** Adam C. English, Fabio Cunial, Ginger A. Metcalf, Richard A. Gibbs, Fritz J. Sedlazeck

## Abstract

Accurately genotyping structural variant (SV) alleles is crucial to genomics research. We present a novel method (kanpig) for genotyping SVs that leverages variant graphs and k-mer vectors to rapidly generate accurate SV genotypes. We benchmark kanpig against the latest SV benchmarks and show single-sample genotyping concordance of 82.1%, which is higher than existing genotypers averaging 66.3%. We explore kanpig’s applicability to multi-sample projects by benchmarking project-level VCFs containing 47 genetically diverse samples and find kanpig accurately genotypes complex loci (e.g. SVs neighboring other SVs), achieving much higher genotyping concordance than other tools. Kanpig requires only 43 seconds to process a single sample’s 20x long-reads and can be run on PacBio or ONT long-reads.

## Introduction

The ever increasing availability of long-read sequencing has begun to enable applications in population scale genomic studies^1,2^. This is possible due to crucial improvements on sample requirements, error rates, and costs for running long-read sequencing instruments. Bioinformatics applications which leverage long-reads are also maturing. These innovations lead to the ability to produce a fully genotyped variant file (VCF) that holds information on each variant’s presence within each sample. This VCF then provides a foundation for subsequent analyses such as genome wide association^3^ and population genomics^4^. While the process of creating fully genotyped VCFs is streamlined for smaller mutations (SNV and indels)^5^ it is not for structural variants (SVs; i.e. genomic alterations larger than 50 base pairs). The current state of the art workflows involve per-sample discovery of SVs which are then merged with methods such as truvari^6^ before subsequently reassessing each SV’s presence or absence (i.e. genotyping) across all samples^1^.

As discovery of SVs using long-reads has improved^2^, genotypers have been introduced which can leverage these SVs. Genotyping is the process of determining the presence of an allele in a sample based on sequencing evidence^7^. This is separate from variant discovery which primarily aims to resolve the structure of alternate alleles. This separation is best illustrated in the case of a heterozygous SV in a diploid organism: After discovery of the alternate allele, a genotyper’s task is to report that both the reference and alternate allele are present in the sample. Therefore, genotypers rely heavily on discovery tools and must be robust to deviations between the reported allele structure and sequencing evidence supporting said allele (e.g. shifted breakpoints)^7^.

State-of-the-art genotypers employ various algorithmic approaches including simple assessments of SVs directly from mapped reads^8^, analysis of variant and read k-mers^9^, and graph-based realignment approaches^10,11^. One thing these approaches have in common is that they have mainly been benchmarked on a single sample. The most commonly used SV benchmark is the GIAB v0.6 benchmark for HG002^12^. This benchmark was built as a consolidation of short-read and noisy long-read discovered variants with Tier 1 regions defined to exclude segmental duplications and complex SVs. However, other SV studies which have used newer long-reads exclusively or whole-genome assemblies have shown that the number and complexity of SVs is greater than the subset found in the GIAB v0.6 Tier1 regions^13^. Furthermore, genotyping SVs across a population increases evaluation complexity by increasing the occurrences of loci containing multiple neighboring and/or overlapping SV alleles. Previous genotypers have some stratification of their benchmarking results based on the presence of neighboring SVs^14^, however this is only in the context of a single sample or trios. The diversity of alleles at any given locus, particularly in tandem repeat loci, is expected to grow as more samples are considered^6,15^.

In this work we describe our software for long-read SV genotyping, kanpig. We showcase kanpig through a comprehensive benchmarking framework, evaluating its accuracy using a diverse set of haplotype-resolved long-read assemblies. We compare kanpig’s genotype accuracy with that of other long-read SV genotypers. In addition to evaluating kanpig using high-confidence, single-sample SVs, we also benchmark against SVs discovered by the commonly used SV caller sniffles and multi-sample SV VCFs. Across all experiments, kanpig consistently produced more accurate genotypes and successfully avoided common errors seen in other genotypers, particularly for neighboring SVs within and across samples.

## Results

### Kanpig Algorithm

Kanpig incorporates four major steps (**Figure 1a**). First, a VCF containing SVs is parsed and SVs within a specified distance threshold of one another are identified as a “neighborhood”. Second, kanpig constructs a variant graph from a neighborhood of SVs with nodes holding SVs and directed edges connecting downstream, non-overlapping SVs. Next, a BAM file is parsed for long-read alignments which span the neighborhood of SVs and pileups within user-defined size boundaries (default 50bp to 10kbp) are generated before reads are clustered to identify up to two haplotypes. Finally, a breadth-first search connecting the variant graph’s source and sink nodes is performed to find the path which maximizes a scoring function with respect to an applied haplotype.

**Figure 1:**
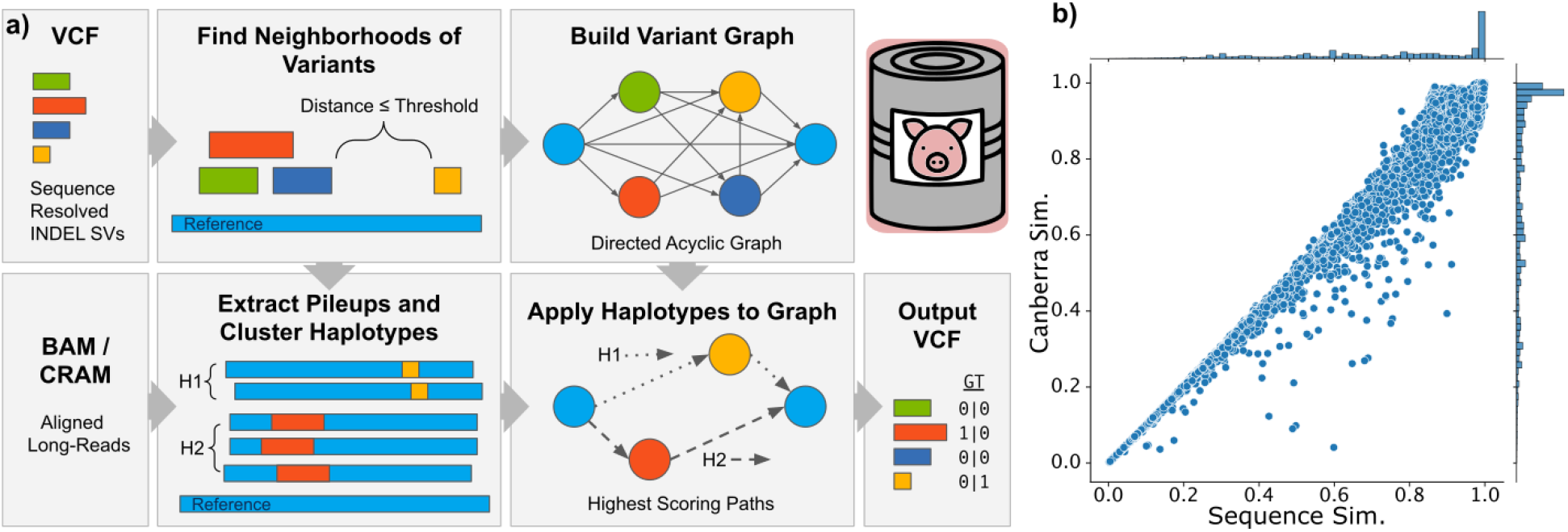
Overview of Kanpig’s Algorithm. a) Schematic of the main steps of kanpig’s algorithm. b) Sequence similarity vs Canberra similarity of ≥50bp insertion pairs within 500bp from GIAB v1.1 SVs.

A novelty of kanpig’s approach is its representation of sequences in a variant graph’s nodes or a read’s pileups as a k-mer vector, using a small k-value, which defaults to 4 base pairs (bp). A k-mer vector contains the counts of all possible k-mers (256 for k=4) and tracks how often each k-mer is observed in a sequence. These vectors are then used as part of the scoring function’s measurement of sequence similarity via Canberra distance^16^. To calculate this similarity, kanpig uses a representative read from a haplotype and includes k-mers that occur a minimum number of times (default: 2) across the read’s pileups and the SV sequences from a path’s nodes. This filtering helps reduce the impact of artifacts, such as sequencing errors. To assess this approach’s accuracy, we compared Canberra similarity (1 minus distance) of k-mer vectors to sequence similarity measured using edlib^17^ over 141,680 insertion SV pairs ≥50bp and within 500bp of one another from GIAB v1.1 SVs^18^. The Canberra similarity of k-mer vectors has a Pearson correlation coefficient of 0.994 (p < 0.01) with traditional sequence similarity (**Figure 1b**) and a mean squared error of 7×10^−4^. Of these nearby insertion pairs, 41,152 (29%) have a size and edlib sequence similarity above 95%, which included 30,641 compound heterozygous pairs occurring at the exact same position. Of these ≥95% similar insertion pairs, 37,918 (92%) also had a Canberra similarity above 95%. Therefore, many neighboring SVs are highly similar and the Canberra similarity metric accurately measures them.

Two crucial components of kanpig’s algorithm are how reads are clustered into haplotypes before applying them to a variant graph and how variant graphs lack edges connecting overlapping SVs. K-means clustering of reads by their k-mer vectors produces up to two haplotypes. Each haplotype then only needs a single search for the optimal path through the graph to apply all its constituent reads simultaneously. Furthermore, by disallowing paths through overlapping variants, kanpig prevents haplotypes from creating conflicting genotypes. For example, in the neighborhood of SVs illustrated in Figure 1a, kanpig ensures a haplotype can only be applied to one of the three overlapping variants. Conversely, if either reads or SVs were evaluated independently, it is possible for reads from the same haplotype to support different, conflicting SVs or for overlapping SVs to redundantly recruit support from the same read. This could give rise to problems such as two overlapping SVs being genotyped as homozygous. Since human autosomal chromosomes are diploid, a homozygous variant which overlaps another present variant in a single sample is conflicting as it is biologically implausible to e.g. lose both copies and then additional copies of overlapped deletions at the same time. Kanpig’s design prevents this type of genotyping error by correctly handling overlapping and nearby SV neighborhoods.

### Establishing a baseline of structural variant genotypes

The Genome in a Bottle consortium (GIAB) has released two sets of structural variants researchers use to evaluate the performance of their SV discovery and genotyping tools. The first, GIAB v0.6^12^, was constructed from a merge of multiple SV discovery tools’ results and the second, draft GIAB v1.1^18^, derived from a single highly polished diploid Telomere-to-Telomere assembly. These two benchmarks of insertions and deletions greater than 50bp have vastly different levels of completeness. GIAB v0.6 contains 9,646 SVs compared to 27,560 SVs for v1.1. The ratio of heterozygous to homozygous variants (het/hom ratio) in GIAB v0.6 is 1.16 compared to 4.15 for v1.1. These benchmarks’ differences are especially notable in the context of overlapping and closely adjacent SVs. While GIAB v0.6 has no Tier1 passing SVs within 1kbp of another SV, 15,685 (43%) of GIAB v1.1 SVs have at least one neighbor. These observations of het/hom ratio and neighboring SVs are not independent as 92.2% of GIAB v1.1 SVs with at least one neighbor are heterozygous compared to 65.5% without neighbors being heterozygous (chi-squared test p<0.01). Therefore, SV genotypers benchmarked primarily against GIAB v0.6 may be biased towards a subset of homozygous SVs.

To comprehensively evaluate kanpig, we leveraged high-confidence assemblies from the Human Pangenome Reference Consortium (HPRC)^19^ to create SVs from 47 genetically diverse genome assemblies using dipcall^20^ alignment to GRCh38 (Methods: Data Collection). To ensure the assembly-derived SVs are of high quality, we compared the HPRC HG002 SVs against GIAB v1.1 SVs using truvari^6^. This resulted in a 0.996 precision and 0.987 recall with a 99.3% genotype concordance. The consistency of the HG002 variants from the HPRC assembly and dipcall pipeline to GIAB v1.1 suggests the pipeline is sufficient for generating baseline VCFs across all HPRC samples.

### Single Sample SV Genotyping

To assess kanpig’s performance, we collected 32x coverage of PacBio long-reads derived from the same 47 HPRC individuals which comprise the assembly-derived baseline variants described above (Methods: Data Collection). We genotyped autosomal SVs between 50bp and 10kbp for each sample using kanpig and three other long-read SV genotypers of SVJedi-graph^14^ (heretofore referred to as SVJedi) and two long-read SV discovery tools with genotyping-only (a.k.a. “force-calling” or ‘-fc’) modes: sniffles^21^, and cuteSV^22^. The average genotyping concordance of a sample’s kanpig predicted genotypes was 82.1% compared to 74.2% for SVJedi, 67.1% for sniffles-fc, and 63.8% for cuteSV-fc. Measurement of genotyper performance was repeated across two down-samplings of the reads to 16x and 8x coverage (**Figure 2a**). It was observed that lower coverage causes lower genotyping concordance. Notably, kanpig’s performance at 8x (77.8%) was higher than other tools’ performance at 32x.

**Figure 2:**
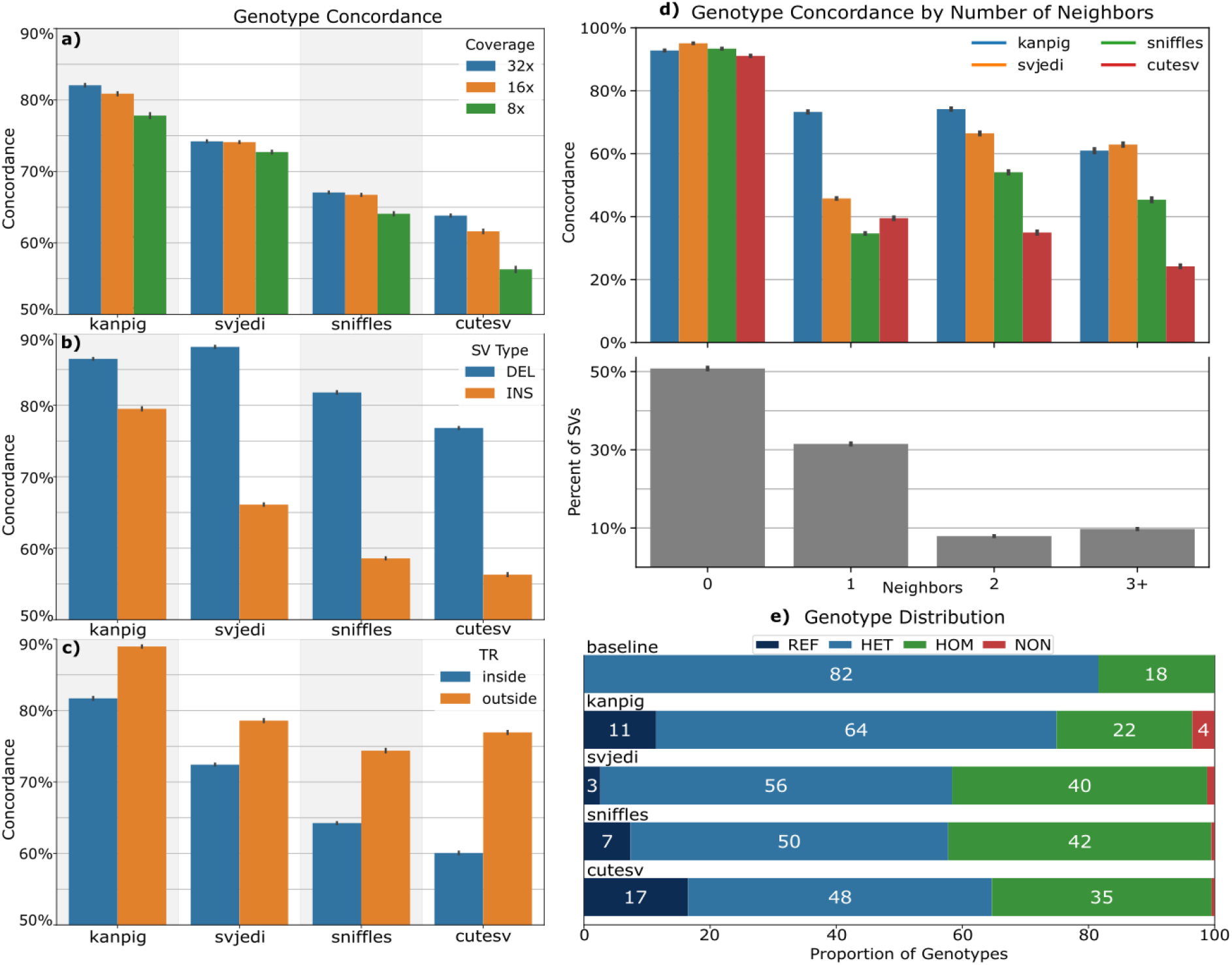
Performance of genotypers on single-sample assembly-derived SVs across 47 samples. Left side: Genotype concordance a) for three read coverages b) stratified by SV type of deletion (DEL) and insertion (INS) c) stratified by SV location inside or outside tandem repeats. Note that left figures have truncated y-axes (50%-90%) for readability. d) Genotype concordance by number of neighboring SVs within one thousand base pairs (top) and proportion of SVs by number of neighbors (bottom). e) Average genotype distribution per sample in the baseline assembly-based variants and for each tool’s result. REF = reference homozygous, HET = heterozygous, HOM = homozygous alternate, NON = Missing genotype (./.). All bar plot error bars represent 95% confidence intervals.

The assemblies produced an average of 9,706 deletions and 16,870 insertions per-sample. Stratifying genotyper performance by SV type found that all tools’ deletion genotypes had higher concordance than their insertion genotypes (**Figure 2b**). The best performing genotyper on deletions was SVJedi at 88.2% average genotype concordance followed by kanpig at 86.5%. For insertions, kanpig’s 79.5% genotype concordance was much higher than SVJedi’s at 66.1%. Kanpig’s performance between SV types had the lowest imbalance (i.e. difference between deletion and insertion concordance) at 7.0 percentage points (p.p) compared to 22.1 p.p. for SVJedi, 23.2 p.p. for sniffles-fc, and 20.5 p.p. for cuteSV-fc.

Another important stratification for SVs is their overlap with tandem repeats since they harbor ∼70% of SVs and pose unique challenges due to their sequence context^15^. We found that kanpig’s genotype concordance on SVs within TRs was 81.7% and 89.0% outside of TRs (**Figure 2c**). SVJedi had the second highest performance at 72.4% inside and 78.6% outside of TRs.

A strong predictor of a genotyper’s ability to correctly assign a genotype was the presence of neighboring variants since they increase the complexity of the locus being evaluated. We measured tools’ genotyping concordance stratified by variants having neighbors within 1,000 bp and those without (i.e. isolated) (**Figure 2d**). In the 49.2% of SVs per-sample with neighbors, kanpig had the highest average genotyping concordance at 70.9%, followed by SVJedi at 52.5%, sniffles-fc at 39.9%, and cuteSV-fc at 35.7%. Over the remaining 50.8% of isolated SVs per-sample, SVJedi had the highest average genotype concordance at 95.0% followed by sniffles-fc at 93.3%, kanpig at 92.7%, and cuteSV-fc at 91.0%. This suggests that all tested genotypers perform reasonably well on isolated SVs while kanpig’s ability to correctly genotype SVs in close proximity to one another differentiates it. The full table of tools’ average genotyping concordance across samples by coverage, SV type, and overlap with TRs is available in Supplementary Table 1.

**Table 1:**
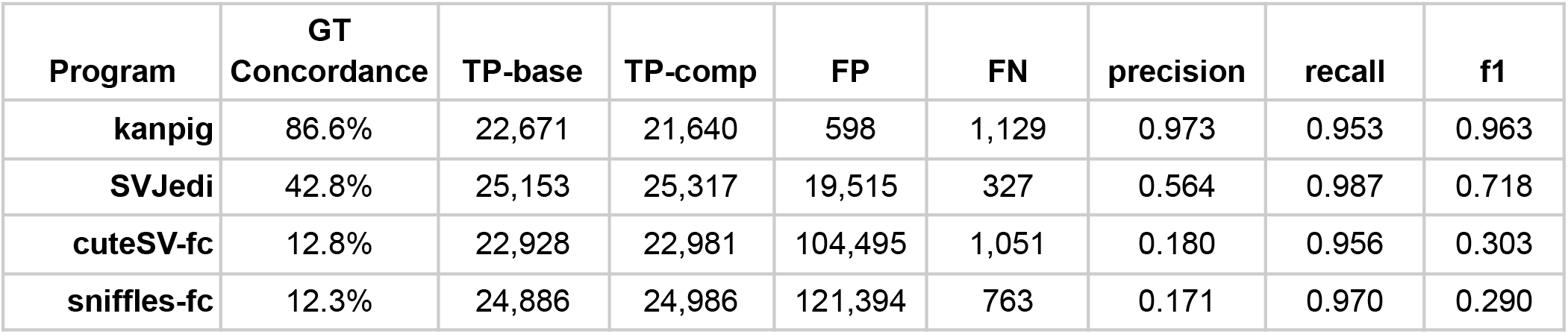
Multi-Sample SV Genotyping Performance. Average performance of genotypers across 47 samples when given a multi-sample VCF of assembly-derived SVs and 32x long-read coverage. SV matching was performed using truvari bench and is detailed in Methods: Measuring Genotyping Performance.

As observed in the GIAB v1.1 SVs described above, there are approximately four times as many heterozygous SVs as there are homozygous SVs expected in an individual, and heterozygous SVs more frequently neighbor another SV than homozygous SVs. We next explored the distributions of genotypes in the assembly-derived SVs to confirm this observation and assess each tool’s ability to classify both heterozygous and homozygous SVs (**Figure 2e**). We first found kanpig’s 3.00 average het/hom ratio per sample is closest to the assembly SVs’ average het/hom ratio of 4.58. The other tools had a stronger bias towards assigning homozygous genotypes with SVJedi having a het/hom ratio of 1.38, sniffles-fc 1.10, and cuteSV-fc 1.38. Since every variant assessed in this experiment is known to be present according to their respective assemblies, any assignment by a genotyper of a reference homozygous state is an error. Kanpig exhibits the second highest average reference homozygous error rate at 11.4% behind cuteSV-fc at 16.5%. However, the above genotype concordance measurements are already penalized for these mistakes and did not prevent kanpig from recording the highest genotyping concordance. Finally, the highest missingness rate (i.e. insufficient evidence to assign any genotype: ./.), is from kanpig with an average of 3.5% of variants per-sample while SVJedi is second at 1.2%.

These measures of genotype concordance, het/hom ratio, reference homozygous errors, and missingness rate suggest that in some stratifications other tools may be able to confirm the presence of alternate alleles (i.e. recall) slightly better than kanpig. However, kanpig’s consistently high assignment of correct genotypes (i.e. precision) across all stratifications lead to a significantly higher overall genotyping accuracy.

### Single Sample Discovery Genotyping

Most projects generating SVs will not have access to high-quality assemblies but will instead discover SVs from raw reads’ alignments using tools such as sniffles^21^. SV discovery tools are susceptible to errors in the form of imperfect variant representations, false negatives, and false positives, each of which challenges the subsequent use of SV genotypers. To test kanpig’s ability to handle errors in the variant graph, we generated SVs on the 47 HPRC 32x samples using sniffles discovery, genotyped with the tools being tested, and compared results to the assembly-derived SV set. On average, kanpig’s average genotype concordance on the 32x coverage experiments was 85.0% compared to 76.2% for SVJedi, and 78.1% for cuteSV-fc. Since sniffles was used to discover the SVs, we measured both its original discovery genotypes as well as its force-genotyping modes’ concordance separately at 80.5% and 80.4%, respectively. In general, the same patterns of genotypers’ relative performance by coverage, SV type, and TR status hold when analyzing the sniffles-discovery VCF as was seen when analyzing the assembly-derived VCFs described above (Supplementary Table 2).

In addition to genotype concordance, we measured how the genotypers changed the precision and recall of present (heterozygous or homozygous alternate) discovered variants. A perfect genotyper would call all false positives (FP) as reference homozygous - thus ‘removing’ the FP - while preserving a present status on all true positives (TP). Unsurprisingly, sniffles force-genotyping on the sniffles discovery SVs created the least amount of change with only 23 FPs and 87 TPs genotyped as reference homozygous per-sample. Kanpig removed an average of 52 FP variants and 206 TPs whereas cuteSV-fc removed an average of 51 FP variants and 745 TP variants. Interestingly, while SVJedi only lost an average of 50 TP variants per-sample, it also found 63 additional FP variants per-sample. These additional false positives were caused by discovered variants which originally had poor quality and a reference homozygous genotype but SVJedi determined the variant was present and therefore created an additional present FP. Supplementary Table 3 contains a full report of tools’ impact on the average precision and recall across the 47 samples and on three read coverages of 32x, 16x, and 8x.

### Computational Performance

Kanpig was written in Rust in order to achieve high speed and memory safety and is available under an open source MIT license. Kanpig can be built from source or run with available statically-linked binaries for POSIX-compliant systems. To test kanpig’s speed and memory usage relative to other genotypers, we first tested each tool’s computational performance on chr1 of a single 12x sample when given only a single core. Of the tools which operate on a BAM file, kanpig was the fastest and least memory intensive at 93 seconds (s) and 17.6 megabytes (Mb) max resident set size followed by cuteSV-fc at 160s (813.9Mb) and sniffles-fc at 350s (209.3Mb). SVJedi’s core code took 7 seconds to run, however it needs unaligned reads in FASTQ format as input in order to run minigraph^23^ read alignment which took 4.6 hours. For context, pbmm2^24^, which was used to create the BAM file given to other genotypers, took 4 hours. All tested genotypers can run using multiple threads and therefore were also tested when given 16 cores and a whole-genome 20x sample. Again, kanpig was the fastest at 43s (1.7 gigabytes (Gb) memory) followed by sniffles-fc (56s / 1.3Gb), and cuteSV-fc (63s / 1.2Gb). SVJedi needed 24Gb of memory and, after discounting the 2h 15m for minigraph runtime, took 95s.

### Multi-Sample SV Genotyping

The input to a genotyper analyzing SVs across multiple samples is a VCF containing SV discovery results per-sample which have been consolidated to remove redundant SVs between samples. To test kanpig’s ability to process a multi-sample VCF, we merged the sniffles discovery variants on the 47 HPRC samples introduced above and used truvari^6^ to collapse putatively redundant variant representations of the same type, within 500bp, and ≥95% similar in sequence and size (Methods: Data Collection). The initial set of discovered SVs had 561,735 total variants and truvari collapse left 257,323 total variants (54.1% reduction) were input into the genotypers. Kanpig had the highest genotype concordance to the baseline SVs at 84.9% followed by SVJedi at 55.4%, cuteSV-fc at 34.1% and sniffles-fc at 33.8%. Furthermore, the programs’ present SV f1 score (harmonic mean of precision and recall) was 0.948 for kanpig, 0.817 for SVJedi, 0.611 for cuteSV-fc, and 0.605 for sniffles-fc. Details of genotyper performance on multi-sample discovered SVs are available in Supplementary Table 4.

While kanpig’s multi-sample performance was consistent with its single sample SV benchmarking, the other tools had a remarkable decrease in genotyping concordance. To ensure that the processes of SV discovery and SV merging were not confounding factors to these changes in performance, we took the assembly-derived SVs per-sample and consolidated them with bcftools merge^25^. This multi-sample input VCF therefore holds the entire set of un-manipulated high-confidence SVs. By not merging the SVs before genotyping, the input VCF contains many highly similar SVs which are entirely valid representations within the context of their derived assembly but ostensibly redundant representations in the context of multiple samples^26^. The consolidated assembly-derived SVs from the 47 HPRC samples contained 403,031 autosomal SVs between 50bp and 10kbp. A total of 371,771 SVs (92.2%) were within 1,000bp of another SV; this included 346,843 (86.0%) within 500bp of another SV of the same type that was also at least 95% similar in terms of size and sequence.

Kanpig had the highest genotype concordance for the assembly-derived, multi-sample SVs at an average of 86.6% across samples followed by SVJedi at 42.8% (**Table 1**). CuteSV-fc and sniffles-fc had an even lower average genotype concordance of 12.7% and 12.2%, respectively. The average kanpig genotyped sample had a precision of present SVs (non-reference homozygous) of 0.973 (mean 598 FPs) while SVJedi had a precision of 0.564 (19,515 FPs). Both cuteSV-fc and sniffles-fc averaged over 100,000 FP SVs per-sample, dropping their precision to under 0.20. Simultaneously, kanpig maintained a reasonable recall of 0.953, which was lower than the other genotypers which achieved a recall of at least 0.956. These observations indicated that kanpig was able to maintain a high genotyping concordance for multi-sample SV genotyping due to having high precision while the other genotypers falsely applied reads to alternate alleles not present in a sample. Details of genotyper performance on assembly-derived multi-sample SVs by coverage, SV type, and overlap with TRs are available in Supplementary Table 5.

For both the sniffles discovery/truvari merge SVs and assembly-derived SVs, the strongest predictor of genotyping concordance was the presence of neighboring SVs (**Figure 3**). On the 32x coverage experiments, SVs without neighbors within 1,000bp had high genotyping concordance between 87.9% (kanpig) and 75.6% (SVJedi) on discovered SVs and between 97.5% (kanpig) and 82.0% (SVJedi) on the assembly-derived SVs. These isolated SVs account for only 16.5% of the discovered SVs and 7.8% of the assembly-derived SVs. For the remaining SVs having at least one neighbor, kanpig’s genotyping concordance was 83.2% on the discovery-derived SVs and 83.8% on the assembly-derived SVs. The second highest performing tool was SVJedi with genotyping concordance of 47.1% and 37.3% for discovered and assembly SVs, respectively. Both sniffles-fc and cuteSV-fc had genotyping concordance below 25% on SVs with neighbors in the multi-sample VCFs.

**Figure 3:**
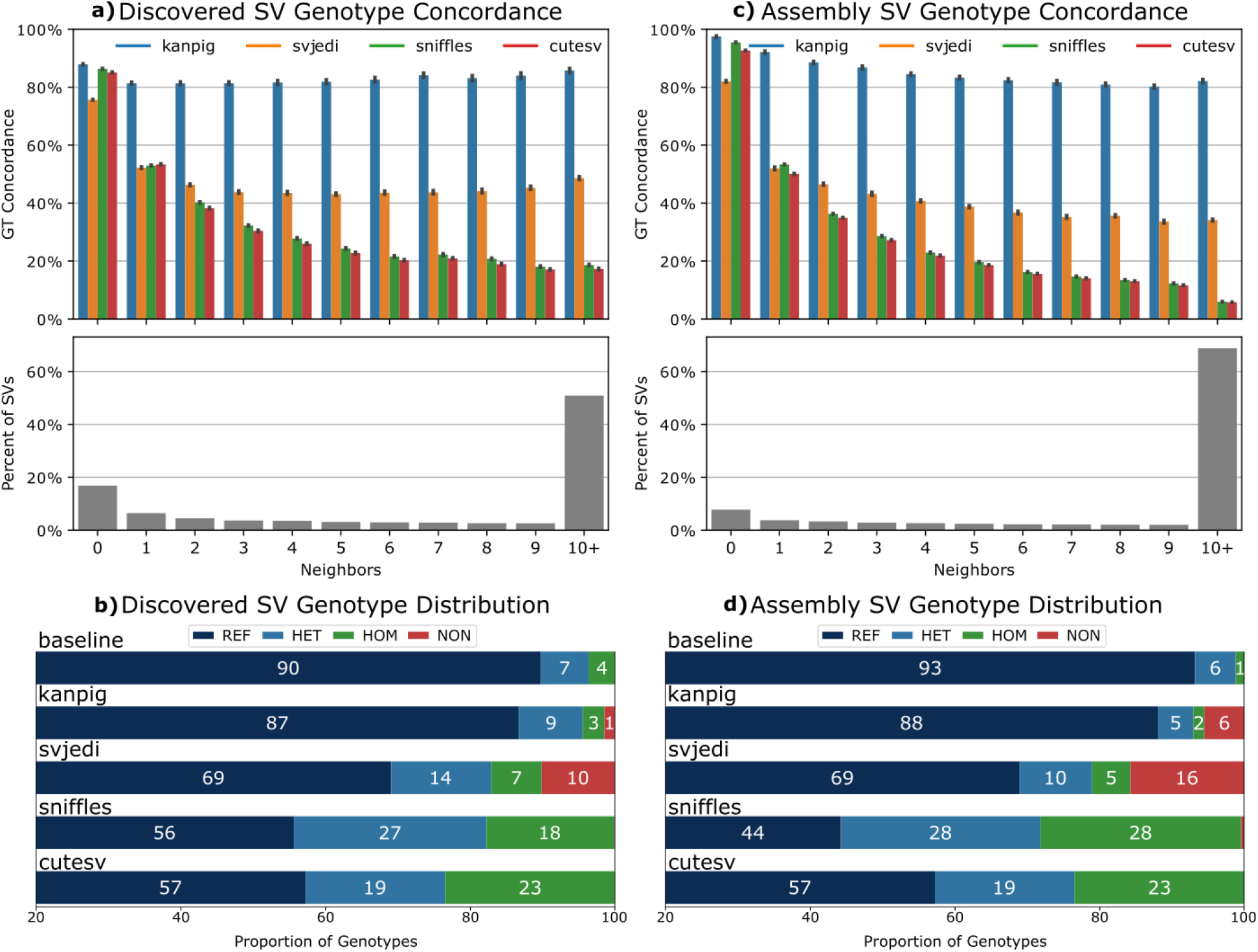
Multi-sample SV genotyping performance. Genotype concordance by number of neighbors for discovered SVs (a) and assembly-derived SVs from 47 samples (b). Genotype distribution for discovered (c) and assembly-derived SVs (d). For b & d, the x-axis has been truncated to 20%-100% for readability. For b, the baseline corresponds to the original sniffles discovery genotypes without full genotyping and all missing (‘./.’) genotypes assumed to be reference homozygous. REF = reference homozygous, HET = heterozygous, HOM = homozygous alternate, NON = Missing genotype (./.). Bar plot error bars represent 95% confidence intervals.

Analysis of the kanpig results found no evidence of conflicting genotypes from homozygous SVs overlapping non-reference-homozygous SVs. However, of the 257k multi-sample discovery SVs, SVJedi had conflicting genotypes from at least one sample for 29.4% of variants, cuteSV-fc had 40.7%, and sniffles-fc had 45.7%. For the multi-sample assembly-derived SVs, the percent of variants with ≥1 samples having a conflicting genotype was again zero from kanpig, 40.5% from SVJedi, 59.1% from cuteSV-fc, and 66.5% from sniffles-fc. The majority of errors from SVJedi, sniffles-fc, and cuteSV-fc were not from mis-assignment between heterozygous or homozygous status but instead an inability to assign a reference-homozygous status to variants which are not present in the sample. To further illustrate where these genotyping errors arise, we investigated the average genotype state distribution per-sample (**Figures 3c/3d**). In the merged assembly-derived VCF, there are 403,031 SVs found across samples. For any one sample, an average of 7% (∼28,000) of these SVs were expected to be present (heterozygous or homozygous alternate). Kanpig genotyped an average of 7% of SVs as present and 6% as missing. SVJedi genotyped 15% of SVs as present and 16% as missing while sniffles-fc and cuteSV-fc genotyped 33% and 66% of all SVs as being present per-sample, respectively, with both producing missingness rates below 1%.

### Performance on Nanopore R9/R10 Sequencing

The above experiments were all performed on PacBio HiFi sequencing, which has a reported read accuracy of ∼99%^27^. Another available long-read sequencing technology is Oxford Nanopore Technologies (ONT) which recently released an update to their sequencing chemistry raising read accuracy from ∼96% (R9 chemistry) to ∼99% (R10 chemistry)^28^. To investigate the ability of kanpig to leverage ONT reads and the impact of read accuracy on genotyping, we ran 3 whole-genome samples with publicly available R9 and R10 replicates having >30x coverage^29^. We found that when genotyping the single-sample assembly SVs, kanpig had a mean genotyping concordance of 77.8% on R9 reads and 80.1% on R10 while SVJedi had 76.9% R9 and 76.1% R10, sniffles-fc 70.8% / 72.3% and cuteSV-fc 70.3% / 71.2%. The comparability of genotyper performance on ONT and PacBio reads suggests all tools can effectively leverage both technologies. However, kanpig’s 2.3 p.p drop in concordance using R9 compared to R10 being higher than other tools (second sniffles-fc at 1.5p.p) indicated kanpig is more dependent on the base-pair level accuracy of reads.

The ONT samples were technical replicates of three of the same individuals in the HPRC PacBio samples. Therefore, we also checked the ability of genotypers to consistently apply genotypes to a sample’s assembly-derived SVs across the replicates. The most consistent genotyper was SVJedi which assigned the correct genotype on all three replicates for 72.2% of SVs. Second was kanpig which had 62.2% consistently correct SVs while sniffles-fc and cuteSV-fc had 55.8% and 49.7%, respectively. When considering only the pair of technologies with higher read accuracy (PacBio HiFi and ONT R10), 72.7% of SVJedi’s genotypes were consistent and concordant, 71.4% of kanpig genotypes, 59.9% for sniffles-fc, and 53.2% for cuteSV-fc (Supplementary Table 6).

## Discussion

Here we presented a detailed description and evaluation of our long-read SV genotyping tool kanpig. By leveraging high-quality SV benchmarks from multiple samples and testing under various conditions, we showed kanpig’s superior accuracy to other genotypers. Our results demonstrated that kanpig consistently outperforms other long-read SV genotypers across most stratifications and use cases. Specifically, we found that kanpig more frequently assigns the correct genotype to nearby (i.e. neighboring) SVs which dominate the SV landscape, particularly when analyzing SVs across multiple samples. In contrast, other widely used methods frequently fail in these situations. An ideal genotyping method for SVs should not just try to maximize the number of SVs present in a sample. This behavior might appear successful for single sample SV genotyping, but, as demonstrated in this work, will lead to issues such as numerous false positives and conflicting genotypes when dealing with a population set of SVs. A major flaw of many SV genotypers is the inability to handle neighboring SVs, which occurs when highly similar, but still distinct SVs occur in close proximity to one another. Neighboring SVs predominantly reside within tandem repeats, which are highly polymorphic and contain around 70% of all SVs in the human genome^15^. If the evaluation of neighboring SVs is improper, genotypes which produce biologically implausible haplotypes per sample (e.g. three overlapping deletions in a diploid genome), can be generated, undermining downstream analyses. Kanpig is able to properly genotype neighboring SVs due to its algorithm evaluating variant graphs constructed over windows of the genome instead of independently searching for read evidence supporting each SV. Furthermore, kanpig’s measurement of sequence similarity between long-reads and paths through the variant graph helps ensure that the single, highest scoring, optimal path through multiple SVs is chosen.

Part of the reason the limitations of other SV genotypers went unnoticed is due to the choice of baseline SVs against which the tools were benchmarked. The GIAB v0.6 SV benchmark has been instrumental in advancing SV discovery algorithms. However, contemporary understanding of SV prevalence and composition highlights the need for more sophisticated benchmarks such as GIAB v1.1 SVs and the HPRC assemblies^19^. As we showed, this benchmark has ∼2.5x more SVs, including ∼43% within 1,000 bp of another SV, and a het/hom ratio of ∼4. Additionally, the HPRC assemblies - which comprise 47 genetically diverse individuals - supported the increased variant count of ∼26 thousand SVs between 50bp-10kbp and ∼4 het/hom ratio per-sample. Not only was kanpig able to achieve the best genotyping concordance on these high-confidence benchmarks, but we also showed kanpig’s ability to accurately genotype discovered SVs from sniffles. While this manuscript only reports test results using sniffles discovery VCFs, kanpig is able to process any compliant VCF with SVs having accurate breakpoints and full allele sequences (i.e. non-symbolic alleles), which are routinely produced by many long-read SV discovery tools.

We benchmarked kanpig’s accuracy when given PacBio HiFi and ONT R9 and R10 to demonstrate its utility across multiple long-read technologies. However, SV genotypers are not only reliant on sequencing technology but also the quality of the provided input SV’s description since genotypers are designed to investigate the postulated SVs. Interestingly, kanpig’s precision in applying sequencing to SVs’ variant graphs was able to reduce the number of falsely discovered SVs while maintaining a high recall, thus, improving not only the calls’ genotypes but also the set of SVs reported per-sample. This is dependent on the information a SV caller provides in terms of accurate and complete allele descriptions. In addition to sequencing technology and SV discovery tools, SV genotyping of multi-sample projects such as pedigrees or populations is also dependent on the SV merging strategy employed. We previously outlined the causes and consequences of over and under merging SVs and implemented our findings in truvari^6^. Alternative methods of SV merging can be utilized to create inputs to kanpig, but researchers need to be aware that over or under merging will impact the performance of not only the SV genotyping methodology but all subsequent analysis.

Any bioinformatics tool designed to assist genomics researchers must undergo continuous development and maintenance. While the version of kanpig presented here can immediately begin serving researchers, future development can implement further improvements. Currently, kanpig is limited to parsing continuous alignments and does not leverage the split or clipped alignment signals that typically occur around larger SVs. As a result, kanpig is currently applicable to variants smaller than the read length. Future versions of kanpig should be able to incorporate split/clipped alignments to better genotype SVs larger than ∼10kbp. Furthermore, kanpig currently requires long-read alignments to span the neighborhood of variants in each locus’s variant graph. However, a future implementation of kanpig could potentially relax this requirement by applying partial alignments to their corresponding subgraph, thereby increasing kanpig’s sensitivity, particularly in low-coverage scenarios. Though this will require careful design to ensure there is no regression in kanpig’s accuracy when genotyping neighboring SVs.

In conclusion, we have described kanpig, a novel SV genotyper, and the potential impacts of improved SV genotyping to SV research. The results presented here on high-confidence benchmarks and other datasets should promote rapid adoption of kanpig for at-scale SV studies, thus improving the reliability of population SV results and potentially contributing to novel findings in biological and clinical research.

## Methods

### Kanpig Algorithm

#### K-mer Vectors

Let `n` be the set of nucleotides `n = {A, G, C, T}` and the function `E` be the two-bit encoding of each element of `n` such that `A=0; G=1; C=2; T=3`. Let `S` be a string of nucleotides with length `l` such that S = n_0_, n_1,_ n_2, …_n_l_. A k-mer `K` of size `k` is a substring of `S` beginning at 0-based position `i` and defined as K_ki_(S) = n_i_, n_i+1_,..n_i+k-1_. K-mer vectors (`V)` hold the counts of all `K(S)`. Each possible k-mer combination of `n` can map to a unique index `p` of a conceptual array `V` with size |n|^k^. The index `p` of K_k0_ can be calculated using a left bitshift operation (`<<`) and the formula:

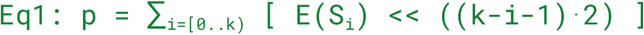

As an example, `p` of the first 3-mer for the sequence “CAGT” is:

`(2 << (2·2)) + (0 << (1·2)) + (1 << (0·2)) = binary 100001 = 33`

V_p_ is then incremented. Next, for every `n` from S_k..l_, we alter `p` via

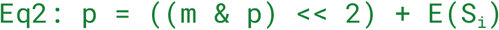

This operation updates `p` by first masking (`m`) the two leftmost bits, left-shifting the remaining bits twice, and adding the current nucleotide’s encoding before again incrementing V_p_. For the example sequence “CAGT”, the second 3-mer’s `p` is 7 (binary 000111).

#### Vector comparison

The similarity of two k-mer vectors can be calculated using the Canberra distance function

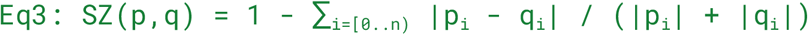

To limit the impact of small sequencing errors, infrequent k-mers (|p_i_| + |q_i_| < minkfreq) are excluded from Eq3. By default, kanpig sets minkfreq to 1, but this threshold can be set by users via a parameter.

In addition to comparing the k-mer vectors’ sequence similarity, the size similarity of two vectors associated with insertion or deletion variants can be compared by separately recording the length of their variants. Deletions constitute a removal of sequence and are thus negative in length whereas insertions add sequence and are thus positive in length. Two vectors’ associated lengths must be of the same sign (e.g. two deletions would both have negative size) to be compared and their size similarity is calculated as

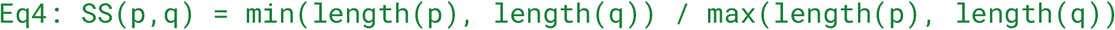

The thresholds for minimum size and sequence similarity needed for two vectors to be considered defaults to 0.90 and can be set by users via a parameter..

#### Variant Graph Construction

A variant call format file (VCF) containing full alleles (REF and ALT sequence) is parsed and partitioned into chunks of variants which are within `--chunksize` distance from one another. This creates sets of variants where the distance between the upstream-most variant’s start position and the downstream-most variant’s end position is at least `--chunksize` base pairs away from any other variant on the same chromosome. A directed graph is constructed with nodes representing each variant and edges linking variants to all non-overlapping downstream variants. A source node upstream of all variants is added to the graph with edges to all variants as well as a sink node downstream with edges from all variants. During conversion of variants into nodes, the reference and alternate sequences are stored as a k-mer vector `V(ALT)-V(REF)`. Additionally, the length of the variant’s described change is recorded as `length(ALT)-length(REF)`.

#### Pileup Generation

For each independent variant graph, read alignments from a BAM file over the graph’s region are parsed. Primary alignments which fully span the region and have above a minimum mapping quality are then piled up to identify insertion and deletion variants within a parameterized size range. Both minimum mapping quality and minimum/maximum size are user parameters with defaults of 5 and 20/10,000, respectively. Each variant’s sequence is converted into a k-mer vector and size is recorded. Reads with identical sets of pileup variants are consolidated and total coverage recorded.

#### Haplotype Creation

The set of pileup variants for each read is then summed to create all putative haplotypes’ k-mer vectors as well as the total size of changes relative to the reference. Putative haplotypes are then grouped with k-means clustering (using k=2) on the k-mer vectors to identify up to two alternate allele haplotypes. For each resultant cluster, the highest covered alternate allele is chosen as the representative for the haplotype. If only one alternate haplotype is identified, the proportions of alternate and reference homozygous read coverage are analyzed (see Methods: Genotyping and Annotation) to identify if the cluster is homozygous alternate or heterozygous. If two clusters are identified, kanpig first compares the size similarity of the two alternate haplotypes and if they are above a threshold ‘hapsim’ they are consolidated and het/hom-alt genotyping performed. Otherwise, the two alternate haplotypes are considered a compound heterozygous pair of alleles and returned to be applied to the variant graph.

#### Applying a Haplotype to a Variant Graph

Given a haplotype and variant graph from a region of the reference, kanpig searches for the path through the variant graph that maximizes the score of the haplotype and path’s similarity. First, a breadth first search starts at the source node and traverses each outgoing edge to create paths to the neighboring nodes. These in-progress paths are pushed to a binary min-heap which prioritizes in-progress paths with lengths more similar to the target haplotype’s size. Kanpig then iteratively analyzes the highest priority paths with the same breadth first search by creating new paths which are extensions of the current path with the head node’s outgoing neighbors. When an edge points to the sink node, kanpig then sums the paths’ variant nodes k-mer vectors and size. Only paths with size and sequence similarity above the user provided minimum thresholds are kept. To allow for false-negatives in the variant graph, when comparing a path to the haplotype, kanpig allows pileup variants in the haplotype to be skipped when summing haplotype k-mer vectors and size (default maximum 3). To account for split-variant representations in the scoring, e.g. the read evidence in the haplotypes suggests two 50bp deletions while the variant graph holds a 100bp deletion. A penalty is incurred for every non-1-to-1 pileup to variant. The algorithm ends its search when either all possible paths have been checked or a user-defined parameter of maximum number of complete paths (default 1000) have been checked.

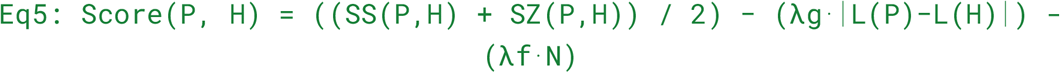

The kanpig scoring function for a path and haplotype is in Eq5 where `P` and `H` are the path and haplotype, respectively, `SS` is the sequence similarity defined in Eq3, `SZ` is the size similarity defined in Eq4. The `L` function measures the length in nodes of a path or number of pileups which comprise a haplotype. The variable `N` is the number of false-negative (i.e. skipped) pileups. The penalty `λg` for split-variant representations defaults to 0.01 while the penalty `λf` for false-negatives defaults to 0.10 and both can be set with user-defined parameters.

#### Genotyping and Annotations

For each variant, the above processes have collected the count of reads supporting the presence of a variant’s alternate allele. The number of reads not supporting the alternate allele’s presence (either reads supporting a second alternate allele or the reference allele) have also been collected. The ratios of reads supporting the presence and absence of a variant’s alternate are then analyzed to determine the variant’s diploid genotype using the same bayesian formula as Chian et.al. ^8^. This formula leverages three priors of allele coverage ratios for reference homozygous, heterozygous, and homozygous alternate read proportions. Kanpig sets these priors to 1e-3, 0.55, and 0.95 for loci with less than 10x coverage and 1e-3, 0.50, 0.90 for loci with at least 10x coverage. The conditional probability of each genotype state given the priors is then calculated and the most likely state assigned. These derived likelihoods are then leveraged to calculate a genotype quality score (confidence in the assigned genotype state) and sample quality score (confidence that the alternate allele is present in any genotype state). Genotype quality score is derived as -10 multiplied by the absolute difference of the second most likely genotype state and most likely genotype state. Sample quality score is derived as -10 multiplied by the absolute difference of the reference homozygous likelihood and the log10 sum of the heterozygous and homozygous alternate likelihoods. Both genotype and sample quality scores range from 0 (lowest quality) to an artificial cap of 100 (highest quality).

In addition to the genotype, genotype quality score, and sample quality score, kanpig will record additional annotations to each variant. These include two common genotyping annotations of depth (DP: total number of reads from a locus analyzed to evaluate this variant) and alternate depth (AD: how many reads from DP support the reference allele and alternate allele). The score described in Eq5 of the haplotypes created for a locus to the final chosen paths through the variant graph are reported as well as a phase-set (PS) unique identifier which annotates the set of variants that comprise each independent variant graph.

Finally, a filter flag (FT) is populated as a bit-flag with states to assist users in filtering genotypes. A flag of 0 can be considered passing. The bits are defined as: 0×1 - The genotype observed from variants matching paths is not equal to the genotype observed from measuring the proportions of reads supporting the two alleles; 0×2 - the genotype quality score is less than 5; 0×4 - the observed coverage is less than 5; 0×8 - the sample quality score is below five for a non-reference-homozygous variants; 0×16 - the alternate allele supporting coverage is below 5; 0×32 - the highest scoring path didn’t use the entire haplotype, which may suggest a false-negative in the variant graph relative to the read evidence.

### Data Collection

HPRC assemblies and PacBio CCS reads were downloaded from the HPRC AnVIL workspace [A]. Only samples in the `assembly_sample` table were considered. CCS reads were downloaded from column `hifi` of the `sample` table, after removing duplicated entries among the BAM and FASTQ files in each cell. For each sample, reads were subset to a desired coverage as follows. Given a list of source files, the list was randomly permuted. For each file in the permuted order, the file was converted to FASTQ if it was a BAM (using `samtools fastq`), and it was appended to a growing FASTQ, until a desired total number of bases was reached or exceeded (measured with `seqkit stats`). Then the FASTQ was randomly permuted (using `terashuf` [B]), and the prefix that achieved the desired number of bases was kept. ONT reads were downloaded from a CARD Google Cloud Bucket [E]. Specifically, we downloaded the `fastq.gz` files from directories `X/reads/` (R9 reads) and `X_R10/reads/` (R10 reads) for every sample `X`, and we downsampled such files as described above.

Given each high-quality assembly and the GRCh38 reference, dipcall was run with default parameters, and multiallelic records were removed from its output using `bcftools norm --multiallelics -`. Finally, only INS and DEL events of length at least 50 bp and with a FILTER field equal to `PASS` or . were kept. To assess the expected quality of the assembly-derived SVs, truvari bench and then truvari refine was run to compare the HG002 assembly SVs to GIAB v1.1 ^18^ using default parameters and a bed file that was the intersection of the GIAB v1.1 high-confidence regions and the dipcall high-confidence regions. The multi-sample dipcall VCF was created by performing a `bcftools merge` of all the single-sample dipcall VCFs, followed by a fina`l bcftools norm --multiallelics -` to remove any remaining multiallelic records.

PacBio CCS reads at each coverage were mapped to GRCh38 with `pbmm2 align --preset CCS` using pbmm2 1.13.0 [C]. Single-sample discovery VCFs were created by running Sniffles 2.3.3 [I] in discovery mode on the 32x CCS BAM of each sample, using default parameters and the tandem repeat track from [D]. The resulting VCFs were then cleaned as follows. BND and CNV records were removed, as well as symbolic INS. The ALT sequence of every symbolic DEL and INV was filled in using the reference, and every DUP was converted into an INS located at the same start position. SVLEN fields were made consistent with REF and ALT fields, and calls longer than 1Mb were removed. The multi-sample discovery VCF was created by running `bcftools merge --merge none` over all the cleaned single-sample VCFs, followed by the following truvari v4.2.2 command:

`truvari collapse --sizemin 0 --sizemax 1000000 --keep common --gt all`

ONT reads were mapped to GRCh38 using minimap2 [F] v2.28 with parameters `-ayYL --MD --eqx --cs`, and specifying `-x map-ont` for R9 reads and `-x lr:hq` for R10 reads.

Long-read genotypers were run as follows. Sniffles 2.3.3 [I] was run in genotyping mode (`--genotype-vcf`) with default parameters. cuteSV 2.1.1 [G] was run with the following flags, derived from its command-line help. For CCS reads:

~~~
--max_cluster_bias_INS 1000 --diff_ratio_merging_INS 0.9
--max_cluster_bias_DEL 1000 --diff_ratio_merging_DEL 0.5
--merge_ins_threshold 500 --merge_del_threshold 500 --min_support
1
~~~

For ONT reads:

~~~
--max_cluster_bias_INS 100 --diff_ratio_merging_INS 0.3
--max_cluster_bias_DEL 100 --diff_ratio_merging_DEL 0.3
--merge_ins_threshold 500 --merge_del_threshold 500
~~~

SVJedi-graph 1.2.1 [H] was run with `--minsupport 1` and all other parameters set to default. Kanpig 0.3.1 was run with a GRCh38 ploidy BED file and with the following parameters for single-sample VCFs:

~~~
--sizemin 20 --sizemax 10000 --chunksize 1000 --gpenalty 0.02
--hapsim 0.9999 --sizesim 0.90 --seqsim 0.85 --maxpaths 10000
~~~

and with the following parameters for multi-sample VCFs:

~~~
--sizemin 50 --sizemax 10000 --chunksize 500 --gpenalty 0.04 --hapsim 0.97
~~~

We tested LRcaller 1.0 [L] on three pairs of 32x CCS BAM and single-sample discovery VCFs, but it got killed by the OS on a cloud VM with 128GB of RAM, so we excluded it from the analysis.

[A] https://anvil.terra.bio/#workspaces/anvil-datastorage/AnVIL_HPRC/data

[B] https://github.com/alexandres/terashuf

[C] https://github.com/PacificBiosciences/pbmm2

[D] https://github.com/PacificBiosciences/pbsv/tree/master/annotations

[E] https://console.cloud.google.com/storage/browser/fc-46bf051e-aec3-4adb-8178-3c51bc5e64ae;tab=objects?authuser=0&pageState=(%22StorageObjectListTable%22:(%22f%22:%22%255B%255D%22))&pli=1&prefix=&forceOnObjectsSortingFiltering=false

[F] https://github.com/lh3/minimap2

[G] https://github.com/tjiangHIT/cuteSV commit 9baddecb32cfbdfbea95be895d0b4edb6cd17fe2

[H] https://github.com/SandraLouise/SVJedi-graph

[I] https://github.com/fritzsedlazeck/Sniffles

[L] https://github.com/DecodeGenetics/LRcaller

### Measuring Genotyping Performance

For the single-sample benchmarking experiments on the assembly-derived variants, SVs between 50bp and 10,000bp from the original VCF and genotyped VCF were merged using `bcftools merge -m none` before the baseline and predicted genotypes were compared. Phase information was ignored e.g. 1|0 is equal to 0|1. SVs on autosomes and with start and ends within dipcall’s reported high-confidence regions were then counted. SVs genotyped as missing (e.g. ./.) were counted separately from the total number of calls and genotype concordance was calculated as the number of matching genotypes over total number of calls.

For all other benchmarking experiments, the assembly-derived variants were matched to the genotyped variants using truvari v4.2.2 and the parameters `bench --pick ac --no-ref a --sizemin 50 --sizemax 10000` along with dipcall’s high-confidence regions as the `--includebed`. The ‘pick ac’ parameter leverages a call’s genotype to determine in how many matches it is allowed to participate. For example, if a genotyper assigns a homozygous status to a variant, its allele count (ac) is two and can therefore match up to two baseline variants. The ‘no-ref a’ parameter ignores variants with a missing or reference homozygous status in the baseline or comparison VCFs. Every genotyped variant with a match and a truvari annotation of GTMatch=0 had an equal baseline genotype and was classified as having a consistent genotype. Every matched variant but with GTMatch != 0 and every unmatched (i.e. false positive) genotyped variant was classified as having an inconsistent genotype. Genotyping concordance is therefore defined as the percent of present calls (heterozygous or homozygous alternate) from the genotyper with a matching variant and matching genotype in the assembly-derived variants.

Stratification of genotyper performance by overlap with tandem repeats was performed by subsetting the adotto TR catalog v1.2^30^ to TR regions contained entirely within an assembly’s dipcall autosomal high confidence regions and only counting an SV as within TR if its start and end boundaries occur within a single TR region. Stratification of genotyper performance by number of neighbors was performed by running `truvari anno numneigh` which annotates an INFO field ‘NumNeighbors’ indicating the number of SVs within 1,000bp for each variant inside the genotyped VCF.

### Measuring Computational Performance

To measure the single-core performance of each tool, we used as input the subset of a 12x CCS HG002 BAM and the subset of the 32x CCS HG002 discovery VCF that relates to chromosome 1 (we used `samtools view` to subset the BAM, `samtools fastq` to get a corresponding FASTQ for SVJedi-graph, and `bcftools view` to subset the VCF). We used only chromosome 1 as the reference. We set the number of threads to one on the command line of each tool, and we pinned each tool to the fifth logical core of our machine with the `taskset` command. We measured wall clock time and maximum resident set size with `/usr/bin/time` (specifically, we timed a set of five iterations of each program, we divided the runtime measure by five and we used the max RSS measure as it is). For minigraph, we used the “Real time” and “Peak RSS” measures that were printed in output.

To measure performance under a typical workload, we ran each tool on a 20x CCS HG002 BAM and on the 32x CCS HG002 discovery VCF over the entire genome, setting the number of threads to 16 on the command line of each tool, and pinning each tool to logical cores 8-23 (all located on the same processor). We ran SVJedi-graph and pbmm2 just once in this experiment.

All experiments were run on a Ubuntu 20.04.4 LTS server (kernel 5.4.0) with two AMD Epyc 7513 processors (64 physical cores in total), 500 GB of RAM, and an NVMe SSD disk. No other user or major task was active.

## Contributions

AE designed the project and implemented kanpig. AE and FC conducted the analysis. AE, FC, GM, RG, and FS wrote the manuscript.

## Code availability

Kanpig is available under an MIT license at https://github.com/ACEnglish/kanpig. Analysis scripts and pipelines used to create and summarize the results presented here are available in the github repository https://github.com/fabio-cunial/kanpig_experiments/.

## Acknowledgements

The authors would like to thank Jason Chin for fruitful discussions on graph-based algorithms, Yulia Mostovy for early beta-testing of kanpig, Michael Macias for development and support of the noodles-vcf rust crate, Xinchang Zheng for early review of the manuscript, and Nathanael Olson and Justin Zook for early access to the GIAB v1.1 benchmark. This manuscript was supported by the Baylor/JHU award (OT2 OD002751, 3OT2OD002751) and NIH award (1U01HG011758-01).

## References

1. Coster, W. D., Weissensteiner, M. H. & Sedlazeck, F. J. Towards population-scale long-read sequencing. Nat. Rev. Genet. 22, 572–587 (2021).

2. Mahmoud, M. et al. Structural variant calling: the long and the short of it. Genome Biol 20, 246 (2019).

3. Sudlow, C. et al. UK Biobank: An Open Access Resource for Identifying the Causes of a Wide Range of Complex Diseases of Middle and Old Age. PLoS Med. 12, e1001779 (2015).

4. Auton, A. et al. A global reference for human genetic variation. Nature 526, 68–74 (2015).

5. DePristo, M. A. et al. A framework for variation discovery and genotyping using next-generation DNA sequencing data. Nat. Genet. 43, 491–498 (2011).

6. English, A. C., Menon, V. K., Gibbs, R. A., Metcalf, G. A. & Sedlazeck, F. J. Truvari: refined structural variant comparison preserves allelic diversity. Genome Biol. 23, 271 (2022).

7. Chander, V., Gibbs, R. A. & Sedlazeck, F. J. Evaluation of computational genotyping of structural variation for clinical diagnoses. Gigascience 8, giz110 (2019).

8. Chiang, C. et al. SpeedSeq: ultra-fast personal genome analysis and interpretation. Nat. Methods 12, 966–968 (2015).

9. Ebler, J. et al. Pangenome-based genome inference allows efficient and accurate genotyping across a wide spectrum of variant classes. Nat. Genet. 54, 518–525 (2022).

10. Garrison, E. et al. Variation graph toolkit improves read mapping by representing genetic variation in the reference. Nat Biotechnol 36, 875–879 (2018).

11. Chen, S. et al. Paragraph: a graph-based structural variant genotyper for short-read sequence data. Genome Biol 20, 291 (2019).

12. Zook, J. M. et al. A robust benchmark for detection of germline large deletions and insertions. Nat Biotechnol 38, 1347–1355 (2020).

13. Audano, P. A. et al. Characterizing the Major Structural Variant Alleles of the Human Genome. Cell 176, 663-675.e19 (2019).

14. Romain, S. & Lemaitre, C. SVJedi-graph: improving the genotyping of close and overlapping structural variants with long reads using a variation graph. Bioinformatics 39, i270–i278 (2023).

15. English, A. C. et al. Analysis and benchmarking of small and large genomic variants across tandem repeats. Nat. Biotechnol. 1–12 (2024) doi:10.1038/s41587-024-02225-z.

16. Lance, G. N. & Williams, W. T. Computer Programs for Hierarchical Polythetic Classification (“Similarity Analyses”). Comput. J. 9, 60–64 (1966).

17. Šošić, M. & Šikić, M. Edlib: a C/C ++ library for fast, exact sequence alignment using edit distance. Bioinformatics 33, btw753 (2016).

18. Zook, J. M. GIAB HG002 Assembly-Based Small and Structural Variants Draft Benchmark Sets. https://ftp-trace.ncbi.nlm.nih.gov/ReferenceSamples/giab/data/AshkenazimTrio/analysis/NIST_HG002_DraftBenchmark_defrabbV0.018-20240716/NIST_HG002_DraftBenchmark_defrabbV0.018-20240716_README_update.md (2024).

19. Liao, W.-W. et al. A draft human pangenome reference. Nature 617, 312–324 (2023).

20. Li, H. et al. A synthetic-diploid benchmark for accurate variant-calling evaluation. Nat. Methods 15, 595–597 (2018).

21. Smolka, M. et al. Detection of mosaic and population-level structural variants with Sniffles2. Nat. Biotechnol. 1–10 (2024) doi:10.1038/s41587-023-02024-y.

22. Jiang, T. et al. Regenotyping structural variants through an accurate force-calling method. bioRxiv 2022.08.29.505534 (2023) doi:10.1101/2022.08.29.505534.

23. Li, H., Feng, X. & Chu, C. The design and construction of reference pangenome graphs with minigraph. Genome Biol. 21, 265 (2020).

24. Li, H. New strategies to improve minimap2 alignment accuracy. Bioinformatics 37, 4572–4574 (2021).

25. Danecek, P. et al. Twelve years of SAMtools and BCFtools. Gigascience 10, giab008 (2021).

26. Audano, P. A. & Beck, C. R. Small polymorphisms are a source of ancestral bias in structural variant breakpoint placement. Genome Res. gr.278203.123 (2024) doi:10.1101/gr.278203.123.

27. Wenger, A. M. et al. Accurate circular consensus long-read sequencing improves variant detection and assembly of a human genome. Nat. Biotechnol. 37, 1155–1162 (2019).

28. Sereika, M. et al. Oxford Nanopore R10.4 long-read sequencing enables the generation of near-finished bacterial genomes from pure cultures and metagenomes without short-read or reference polishing. Nat. Methods 19, 823–826 (2022).

29. Kolmogorov, M. et al. Scalable Nanopore sequencing of human genomes provides a comprehensive view of haplotype-resolved variation and methylation. Nat. Methods 20, 1483–1492 (2023).

30. English, A. Project Adotto Tandem-Repeat Regions and Annotations. Preprint at https://zenodo.org/records/8387564 (2022).

